# A precision virtual crossmatch decision support system for interpretation of ambiguous molecular HLA typing data

**DOI:** 10.1101/756809

**Authors:** Navchetan Kaur, David Pinelli, Evan P. Kransdorf, Marcelo J Pando, Geoffrey Smith, Cathi L Murphey, Malek Kamoun, Robert A Bray, Anat Tambur, Loren Gragert

## Abstract

Virtual crossmatch (VXM) compares a transplant candidate’s unacceptable antigens to the HLA typing of the donor before an organ offer is accepted and, in selected cases, supplant a prospective physical crossmatch. However, deceased donor typing can be ambiguous, leading to uncertainty in compatibility prediction. We have developed a web application that utilizes ambiguous HLA molecular typing data to assist in VXM assessments. The application compares a candidate’s listed unacceptable antigens to computed probabilities of all possible two-field donor HLA alleles and UNOS antigens. The VIrtual CrossmaTch for mOleculaR HLA typing (VICTOR) tool can be accessed at http://www.transplanttoolbox.org/victor. We reanalyzed historical VXM cases where a transplant center’s manual interpretation of molecular typing results influenced offer evaluation. We found that VICTOR’s automated interpretation of ambiguous donor molecular typing data would influence VXM decisions. Standardized interpretation of molecular typing data, if applied to the match run, could also change which offers are made. HLA typing ambiguity has been an underappreciated source of immunological risk in organ transplantation. The VICTOR tool can serve as a testbed for development of allocation policies with the aim of decreasing offers refused due to HLA incompatibility.

## INTRODUCTION

The virtual crossmatch (VXM) procedure involves comparing the transplant candidate’s unacceptable HLA antibodies to a donor’s HLA typing to determine if an organ offer should be accepted^1,2^. A positive VXM that would prompt a refused offer is indicated by the presence of donor-specific antibodies (DSAs)^3,4^.

The adoption of VXM has been an unequivocal success that should be built upon. The changes to the kidney allocation system (KAS) in December 2014 improved equity by giving higher priority to sensitized patients, including regional and national organ sharing^5^. The number of organs allocated post-KAS based solely on a VXM has increased, especially for highly-sensitized patients receiving non-local offers^6–8^. The aim of the VXM in this scenario is evaluation of organ offers without physical testing prior to organ shipment^9^. The VXM has reduced cold ischemia time as well as costs associated with prospective physical crossmatch testing^10^.

Despite the importance of donor HLA typing in determining transplant histocompatibility, molecular HLA typing data is not fully utilized in the United Network for Organ Sharing (UNOS) UNet match run system. The match run relies on a limited vocabulary of UNOS antigens to represent HLA specificities, in contrast to the more precise nomenclature system based on genomic allele sequences curated in the IPD-IMGT/HLA database^11^. Most unacceptable HLA antigens (UAs) defined by Organ Procurement and Transplantation Network (OPTN)^12^ are based on broad and split serologic antigen equivalents^13^. As a result of advances in HLA antibody testing, some two-field allele specificities were recently added to UNet to facilitate allocation especially to the highly-sensitized candidates^14,15^. Although molecular-based testing methods are mandated for deceased donor HLA typing, UNet cannot precisely represent intermediate resolution molecular typing data^16^. Typically deceased donor typing does not fully resolve to unambiguous two-field IMGT/HLA alleles^17^. Because most deceased donor typing is reported at antigen level, many transplant centers do not select allele-specific UAs, as they would not block offers unless the donor is typed at high resolution. Additionally, allele-specific UAs are not used in determining calculated panel reactive antibody (CPRA) values^18^, as two-field HLA frequencies are not included in the UNOS reference panel^19^.

Ambiguous molecular HLA typing data must be manually interpreted within a one-hour time period prior to offer acceptance by the transplant center. Navigating between the serologic and DNA-based HLA nomenclature systems to perform the VXM can be a laborious process that involves interpreting lists of the donor’s possible HLA alleles and comparing those with interpretations of bead patterns from solid phase assays that represent antibody reactivity for a limited panel of HLA proteins. Despite the success of the VXM, unexpected positive physical crossmatch remains common, especially for candidates that are highly sensitized^20^.

This study aims to illustrate the benefits of capturing molecular typing data directly in UNet and having the match run system perform a standardized interpretation of the primary HLA typing data. Advances in HLA antibody identification has led to the appreciation of the presence of allele-specific UAs which has increased the complexity of the allocation system because now one must consider probabilities of high-resolution donor HLA alleles. We draw inspiration from hematopoietic stem cell transplantation registry matching software, where managing HLA typing ambiguity is a major consideration^21^. As a step towards electronic collection and utilization of molecular typing data in UNet, we have built a web application that cross-references each of candidate UAs to the donor HLA typing, providing probabilities that any the UAs will be DSAs. This study intends to test the hypothesis that computer-assisted interpretation of molecular HLA typing data would reveal underappreciated immunological risks stemming from typing ambiguity.

## METHODS

### Data Sources

#### Mapping tables for current IPD-IMGT/HLA alleles and UNOS antigen equivalents

The mapping tables for current IPD-IMGT/HLA alleles (IPD-IMGT/HLA database release 3.370)^22^ to the corresponding UNOS antigen equivalents and reverse mapping from UNOS antigen equivalents to IPD-IMGT/HLA alleles were obtained from our previously described ALLele-to-ANtigen conversion tool (“ALLAN”)^23^ that maps between antigens and alleles in accordance with OPTN guidelines^24^. These tables facilitate translating HLA typing data between the two nomenclature systems of serology-based antigen versus nucleotide-based allele categories for computing VXM.

#### Current policy utilizing UNOS reference tables for UA equivalencies

The list of incompatible donor antigens given each listed UA for the candidate is specified in the OPTN Policies document (Tables 4-5 through 4-10)^12^. These tables consider relationships between broad and split antigens as well as related antigens. If a UA of A2 is listed, donors with the A2 antigen listed will be blocked from offers, along with donors with any of the two-field allele specificities that have A2 as their parental antigen. Under current policy, if A*02:01 is listed as an UA, then only donors with A*02:01 in their HLA typing will be blocked from offers by the match run. Donors with an antigen-level typing A2 are not blocked by an A*02:01 UA.

#### High-resolution HLA allele frequencies for US populations

Allele frequencies for 26 US populations including 5 broad race/ethnic categories (African American (AFA), Asia/Pacific Islander (API), Caucasian (CAU), Hispanic (HIS), Native American (NAM)) and 21 detailed categories were obtained from a published National Marrow Donor Program (NMDP) dataset^25^. Allele frequencies are used to compute probabilities of all possible DSAs when the donor typing is ambiguous.

### Implementation of the Virtual Crossmatch Algorithm

#### Input Data for the Precision Virtual Crossmatch Tool

Our web tool defines two ways to interpret donor HLA typing: “Current UNOS Match Run Logic” and “Proposed Algorithm for Interpreting Ambiguous HLA Typing”. The “Current UNOS Match Run” branch follows current OPTN policy to compute VXM for HLA typing represented either as UNOS antigen equivalents or high resolution IPD-IMGT/HLA alleles. The “Proposed Algorithm” branch interprets ambiguous HLA typing data represented either as genotype list strings (GL String)^26^, NMDP multiple allele codes (MACs)^27^, or UNOS Antigen equivalents. Each possible two-field allele can be assigned a probability. Required input data to compute VXM includes the donor HLA typing, candidate UAs, and donor race/ethnicity (if HLA typing is ambiguous).

#### UNOS Antigen Typing Data Interpretation

When molecular typing data is unavailable, the tool allows for computation of VXM when donor HLA typing is represented at the antigen level, including Bw4 and Bw6 epitopes, allowing for reanalysis of legacy typing data in UNet under a new interpretation scheme. Any high-resolution IPD-IMGT/HLA alleles may be entered as UAs. The expanded list of donor UNOS antigens and Bw4/Bw6 epitopes is then cross-referenced with candidate’s UAs. If there are any conflicting HLA antigens, the VXM is computed as positive. Thus, under the new proposed algorithm, the donor will be blocked if the imputed probability of high resolution DSA is above the provided threshold. Loci supported include A, B, C, DRB1, DRB3/4/5, DQA1, and DQB1. For direct comparison of the proposed interpretation algorithm with current OPTN policy, we also have created a version of our tool that utilizes the current UA equivalency tables.

#### High Resolution Molecular Typing Data Interpretation

If unambiguous two-field HLA allele typing data is available, all antigen equivalencies and corresponding Bw4/Bw6 epitopes are enumerated following the conversion table developed for ALLAN. Bw4 or Bw6 epitopes are mapped to HLA-B locus alleles that harbor the canonical or noncanonical motifs curated by NMDP^28^. Any IMGT/HLA allele can be entered for both the donor HLA typing and the list of UAs. Any conflicts between the donor’s alleles or corresponding UNOS antigen equivalents and mapped Bw4/Bw6 epitopes with candidate’s UA indicate a positive VXM. When high-resolution typing is provided, the tool follows current OPTN policy. Rather than attempting to assign high-resolution alleles from ambiguous typing data, laboratories should use Ambiguous HLA Typing interpretation tool described below.

#### Ambiguous HLA Typing Interpretation

In order to compute VXM for an ambiguous donor HLA typing, the GL strings and NMDP MACs are first mapped to a list of respective possible UNOS antigens and Bw4/Bw6 epitopes using the ALLAN tables. For each HLA locus, all possible UNOS antigens are listed rather than only the UNOS antigens comprising the most probable genotype. For ambiguous UNOS antigen equivalents, the antigens are mapped to all possible high-resolution IPD-IMGT/HLA alleles and Bw4/Bw6 epitopes. The list of high-resolution alleles, antigens, and the Bw4/Bw6 epitopes for the donor typing are cross-referenced with the candidate’s UAs. Considering the HLA typing ambiguity, there will be a probability of VXM positivity provided for each potential DSA. Donor race/ethnicity is used to select the appropriate NMDP population allele frequency distribution for calculating probabilities. If any of the DSA has a probability of higher than the user-defined threshold, VXM is deemed positive. Loci supported by these functions include A, B, C, DRB1, DRB3/4/5, and DQB1. The DQA1, DPA1, and DPB1 loci are not yet included, as comprehensive allele frequencies are under development.

#### Web Application, Command Line Tool, and Web Services Interfaces

An online web application named VIrtual CrossmaTch for mOleculaR HLA typing data (“VICTOR”) that assists in the VXM procedure was developed using the Python Django web framework^29^ and is available at http://www.transplanttoolbox.org/victor. VICTOR is an open source application distributed under GNU General Public License 3. Web services^30,31^ are available for advanced users for remote scripting and integration into HLA lab systems at http://www.transplanttoolbox.org/victor/services. A command-line based application can be installed as a Python package “transplanttoolbox-victor” from the PyPI repository (the Python Package Index). The URL endpoints and commands for the HLA input data types are listed in Table 1. Documentation for the Python package is available at https://transplanttoolbox-victor.readthedocs.io/. The source code for the tool is available in a public GitHub repository at https://github.com/lgragert/virtual-crossmatch/ and archived in Zenodo at https://zenodo.org/record/1252003.

**Table 1:**
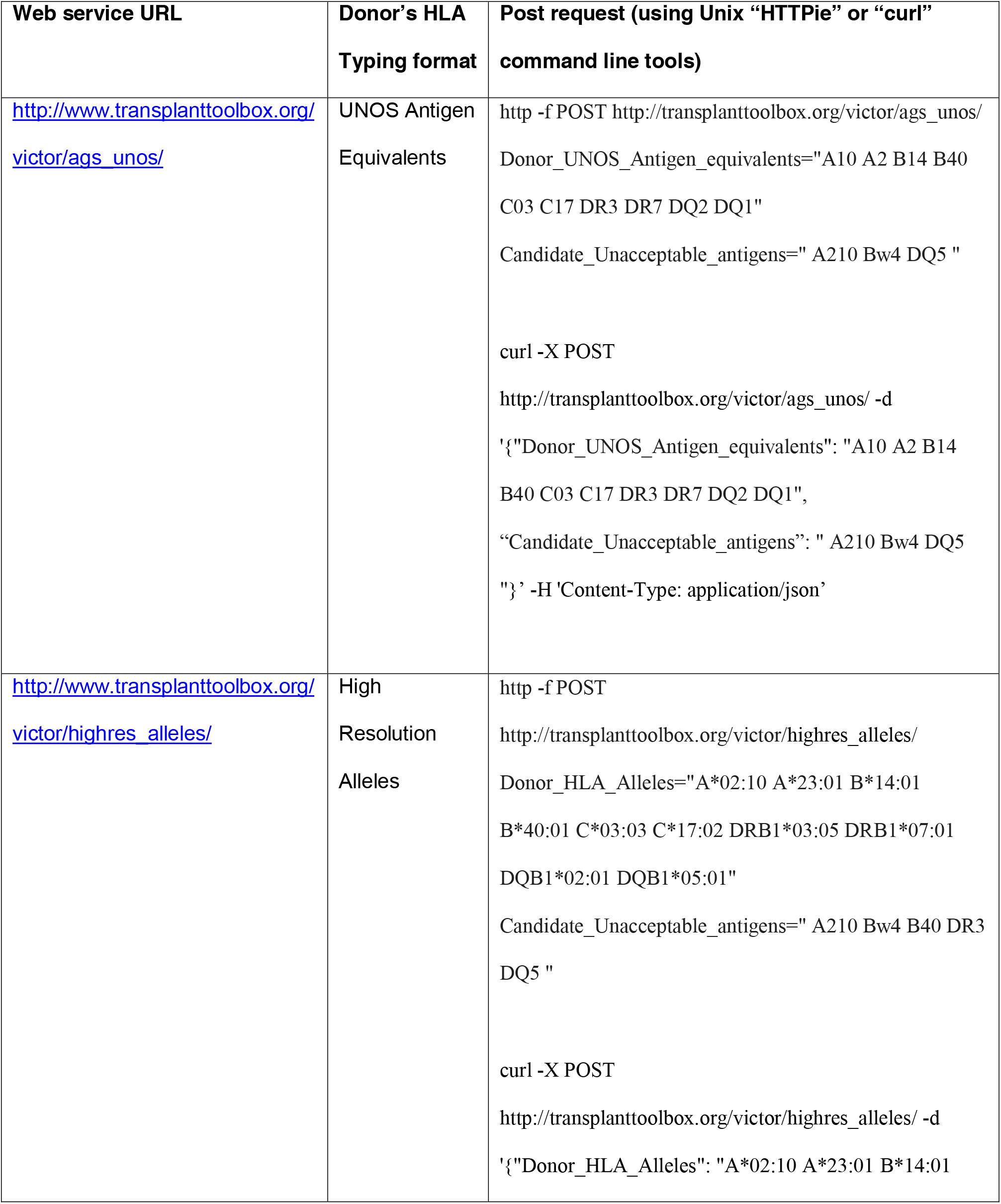
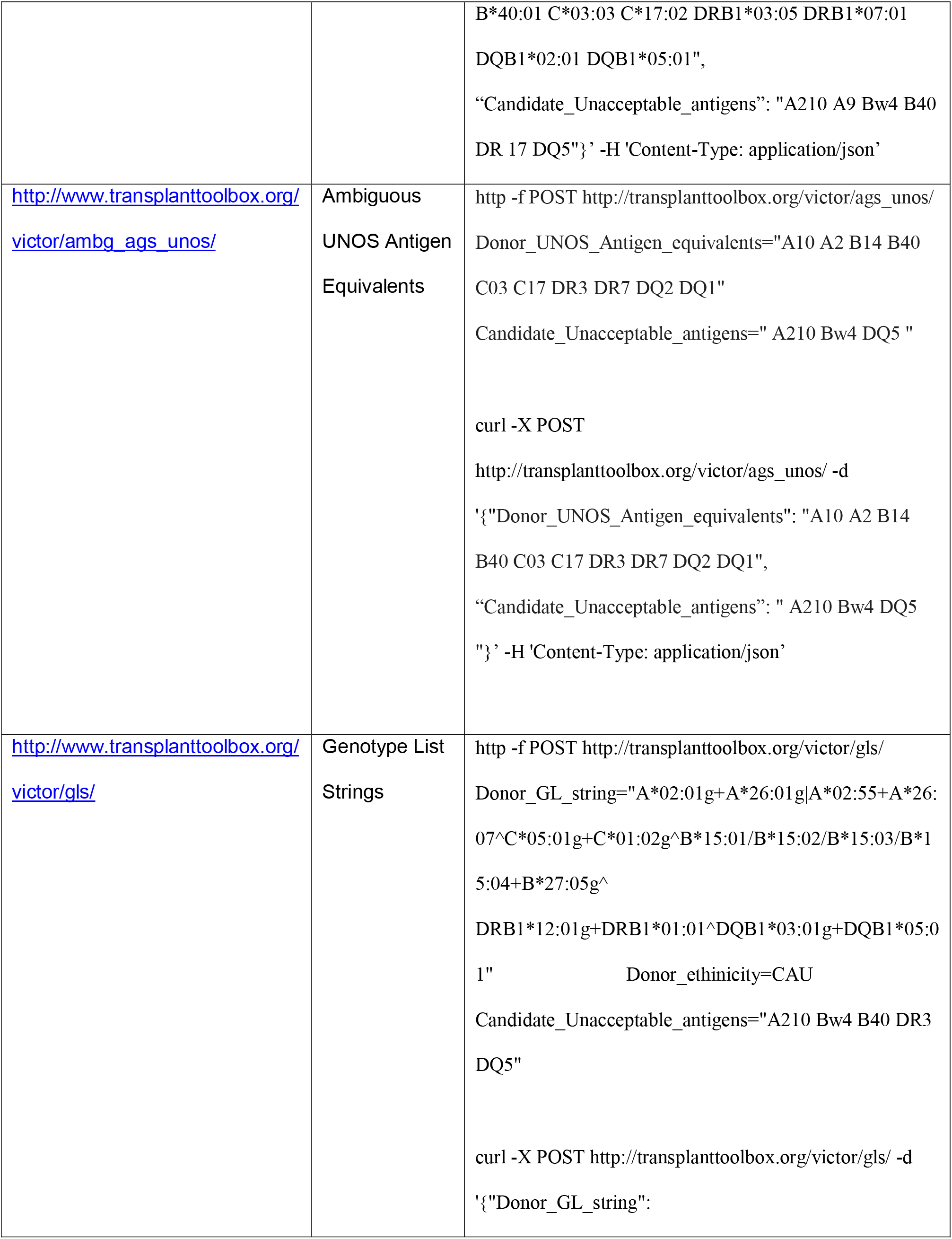
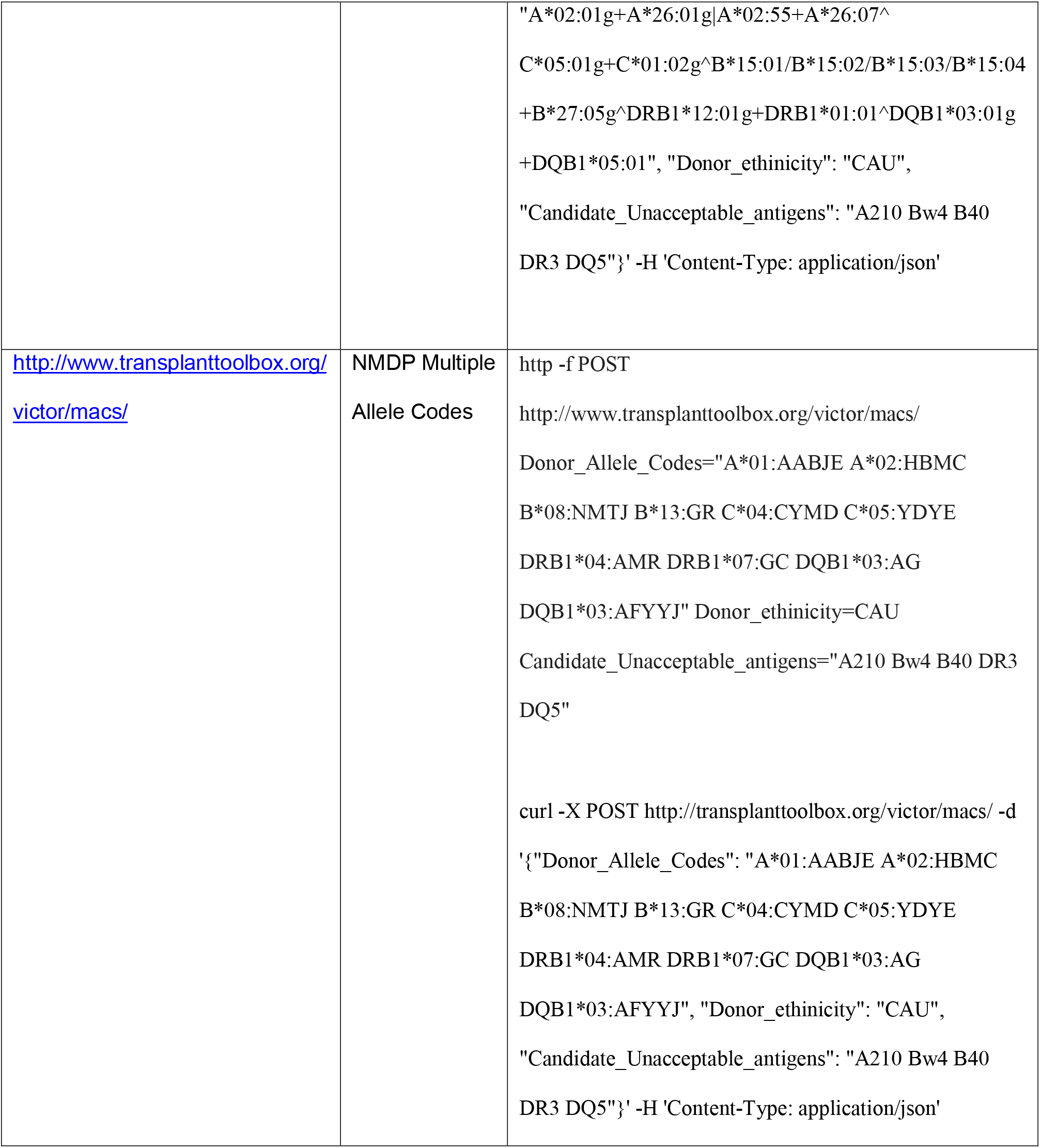
Web services endpoints for VICTOR to compute virtual crossmatch

#### Recomputation of Virtual Crossmatch by VICTOR for Actual Cases of Organ Offers made by UNet

Challenging cases of deceased donor offer evaluations were selected for re-computation of VXM assisted by the VICTOR tool from the following transplant centers: (Northwestern University, Emory University, and University of Pennsylvania). Data from these anonymized cases included the candidate’s UAs and donor antigens entered in UNet as well as donor molecular typing reports attached in UNet or made available from the donor center. This study was approved by the Tulane Institutional Review Board (IRB).

## RESULTS

### Computation of Virtual Crossmatch within a Web Application

Our VXM tool, “VICTOR”, is available as a web application and standalone command line tool. VXM for “Current UNOS Match Run Logic” is computed based on the donor’s HLA typing and the candidate’s UAs. Under the proposed algorithm for interpreting ambiguous HLA typing, four user inputs are required: the donor’s HLA typing, the candidate UAs, the donor’s race/ethnicity, and a probability threshold. The web application is available at http://www.transplanttoolbox.org/victor. The application returns an overall recommendation for VXM along with any DSA probabilities.

### Virtual Crossmatch Case Reports

The analyzed cases were divided into three categories. Category #1 were refused offers where an unlisted allele-specific UA was determined to be high risk. Category #2 were accepted offers where an unlisted allele-specific UA was possible given the donor antigens listed in UNet, but excluded in the molecular typing report. Category #3 were offers were refused due to uncertainty in molecular typing but had low probability of DSAs according to VICTOR. We provide a detailed readout from VICTOR of the probability of positive VXM for each possible DSA for selected cases that illustrate limitations of the current match run system. The categories are summarized in Table 2.

**Table 2:**
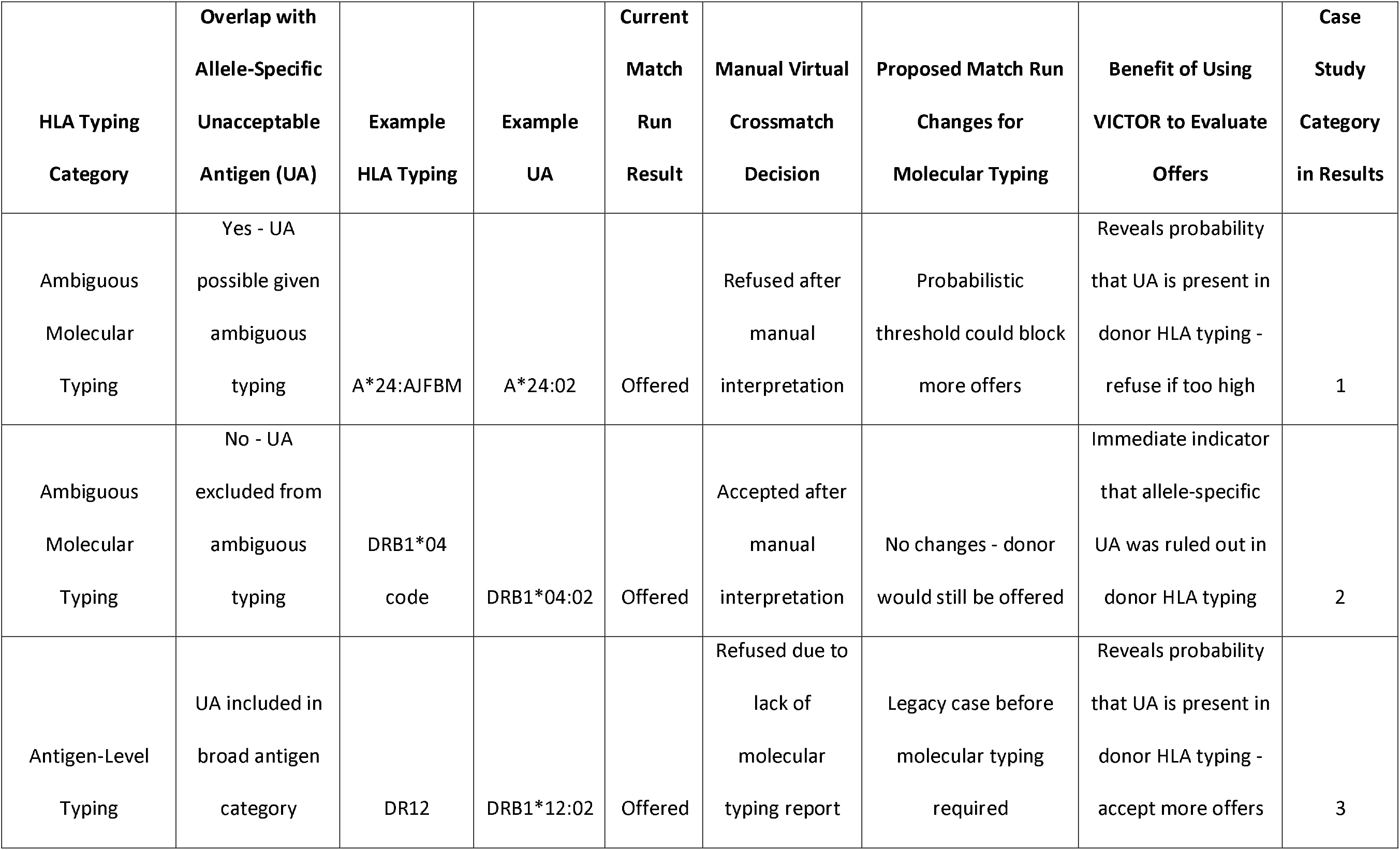
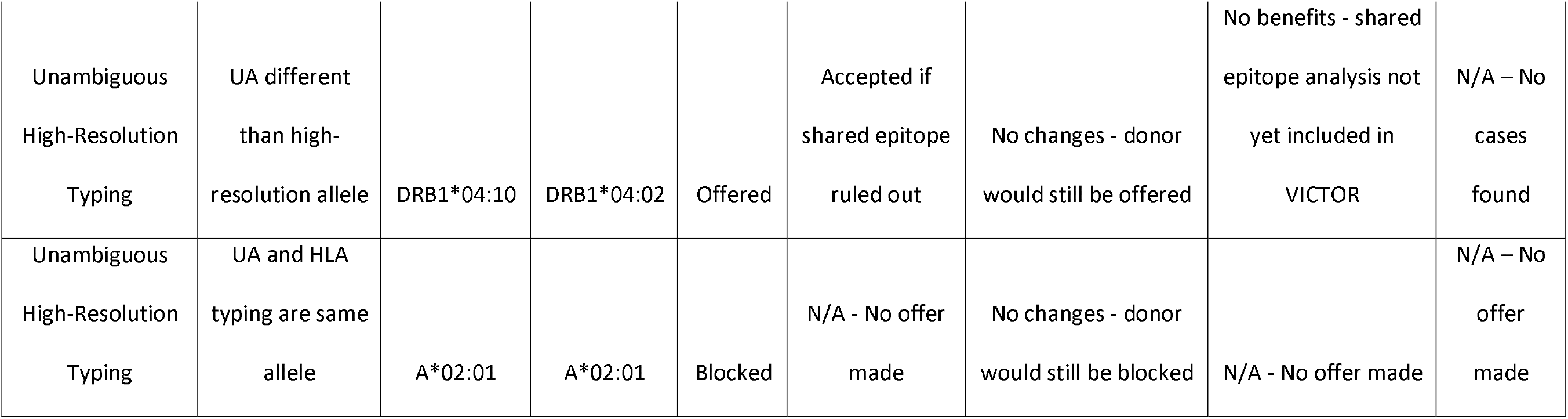
Categories of offers that illustrate how HLA typing ambiguity and allele-specific unacceptable antigens impact match run logic and virtual crossmatch decisions.

| **HLA Typing Category** | **Overlap with Allele-Specific Unacceptable Antigen (UA)** | **Example HLA Typing** | **Example UA** | **Current Match Run Result** | **Manual Virtual Crossmatch Decision** | **Proposed Match Run Changes for Molecular Typing** | **Benefit of Using VICTOR to Evaluate Offers** | **Case Study Category in Results** |
| --- | --- | --- | --- | --- | --- | --- | --- | --- |
| Ambiguous Molecular Typing | Yes - UA possible given ambiguous typing | A*24:AJFBM | A*24:02 | Offered | Refused after manual interpretation | Probabilistic threshold could block more offers | Reveals probability that UA is present in donor HLA typing - refuse if too high | 1 |
| Ambiguous Molecular Typing | No - UA excluded from ambiguous typing | DRB1*04 code | DRB1*04:02 | Offered | Accepted after manual interpretation | No changes - donor would still be offered | Immediate indicator that allele-specific UA was ruled out in donor HLA typing | 2 |
| Antigen-Level Typing | UA included in broad antigen category | DR12 | DRB1*12:02 | Offered | Refused due to lack of molecular typing report | Legacy case before molecular typing required | Reveals probability that UA is present in donor HLA typing - accept more offers | 3 |
| Unambiguous<br>High-Resolution<br>Typing | UA different<br>than high-resolution allele | DRB1*04:10 | DRB1*04:02 | Offered | Accepted if<br>shared epitope<br>ruled out | No changes - donor<br>would still be offered | No benefits - shared<br>epitope analysis not<br>yet included in<br>VICTOR | N/A – No<br>cases<br>found |
| Unambiguous<br>High-Resolution<br>Typing | UA and HLA<br>typing are same<br>allele | A*02:01 | A*02:01 | Blocked | N/A - No offer<br>made | No changes - donor<br>would still be blocked | N/A - No offer made | N/A – No<br>offer<br>made |

### Category #1 Cases: Refused offers due to high risk of donor-specific antibodies

A highly-sensitized patient was listed with the following UAs: A1, A23, A29, A36, A80, B8, B76, DR7, DR17, DR18, DR52. Allele-level UAs for A*24:02 and B*44:02 were identified, but unlisted. The candidate’s antibody assay was negative for A*24:03. A donor with ambiguous typing of A*24:AJFBM was offered, which decodes to a long list of alleles in the A*24 allele family, including A*24:02 and A*24:03. The offer was refused because it was unclear if A*24:02 was present in the donor, and sera was not available to perform a physical crossmatch. The VICTOR algorithm revealed based on allele frequencies that the donor likely had an A*24:02 (probability = 0.94). We also checked the HLA typing using HaploStats, which is based on haplotype frequencies but is unable to interpret some HLA typing with rare alleles. HaploStats and VICTOR were in close agreement. Figure 1 illustrates the computation of VXM by VICTOR for this case. If a VICTOR-based match run had a positive crossmatch probability threshold for making organ offers, this offer would likely not have been made at all, which could have reduced cold ischemia time.

**Figure 1:**
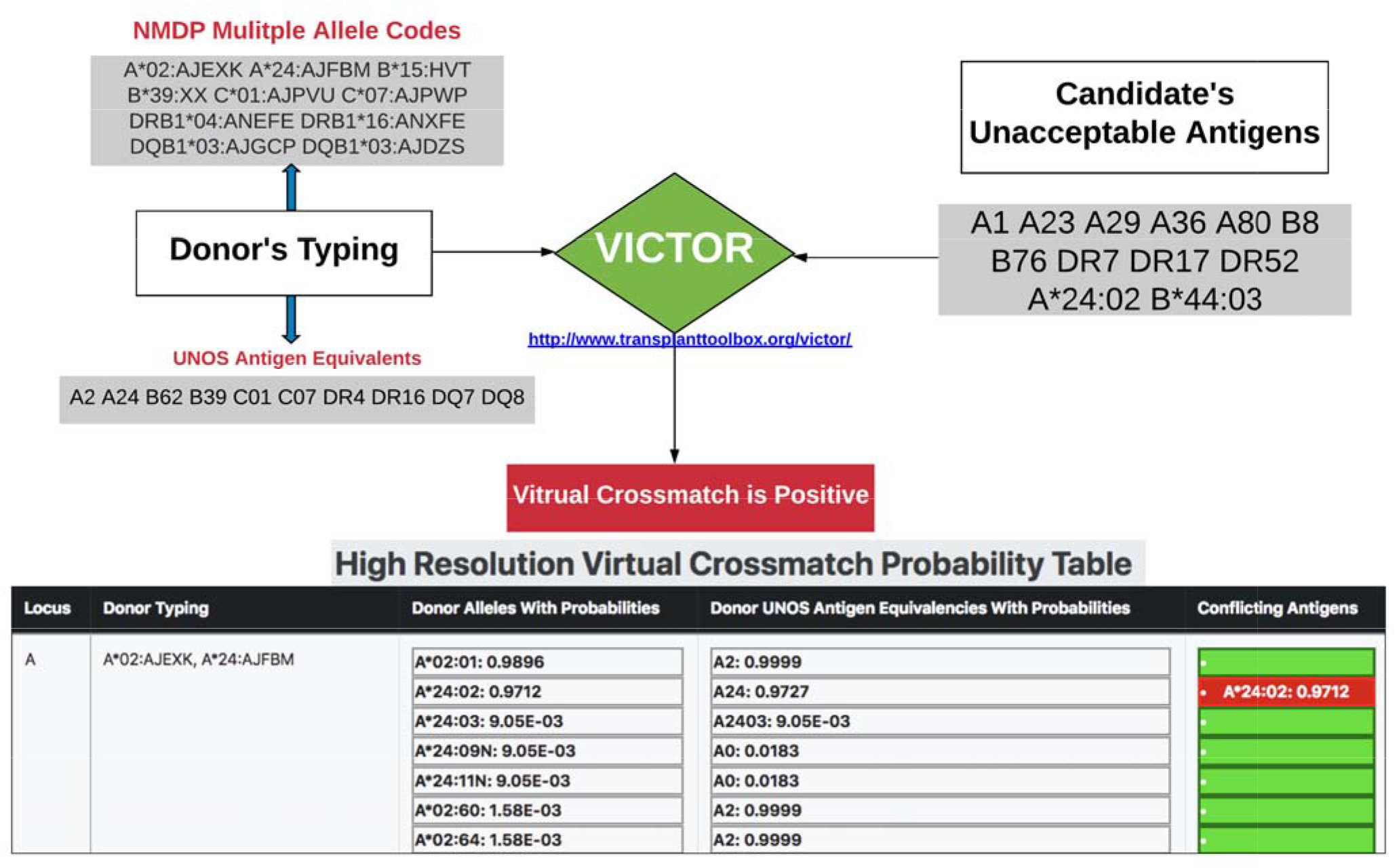
Illustration of a case where virtual crossmatch was computed positive by VICTOR due to high risk of DSA

A second case was similarly refused with a candidate UA of A*02:01 and a donor typed as A2 also returned a risk probability of 0.94 for positive VXM.

A third case was analyzed with candidate UAs of DRB3*02:02 and DRB1*13:03 and donor typing of DR52 and DR13. VICTOR predicted the risk probability of 0.98 for DRB3*02:02 and 0.19 for DRB1*13:03.

### Category #2 Cases: Accepted offers where allele-specific UAs overlapped with the donor HLA antigen listed in UNet, but were ruled out in molecular typing report

A transplant candidate was listed with UAs of Bw4 and DR53. The transplant center also identified an allele-specific UA to A*11:02, but did not list it. The antibody assay had a negative result for A*11:01. In order to receive offers from donors who likely had A*11:01, but were listed with ambiguous HLA typing, A11 was not listed as an UA. The transplant center received an offer from a donor listed with A11 in UNet. Detailed manual examination of the ambiguous molecular typing data revealed that A*11:02 had been ruled out by the typing assay performed. The only possible two-field alleles in the common and well-documented (CWD) allele list were A*11:01 and A*11:04. The offer was accepted. Interpretation of the molecular typing with the VICTOR algorithm also revealed no DSA. The probabilities for the A*11 group alleles in the donor are shown in Figure 2. Here we assumed that the transplant center would have listed any other alleles that shared the putative antibody-reactive epitope with A*11:02 as UAs. We also consider the possibility that the A*11:02 bead could have been a false positive due to a cryptic epitope.

**Figure 2:**
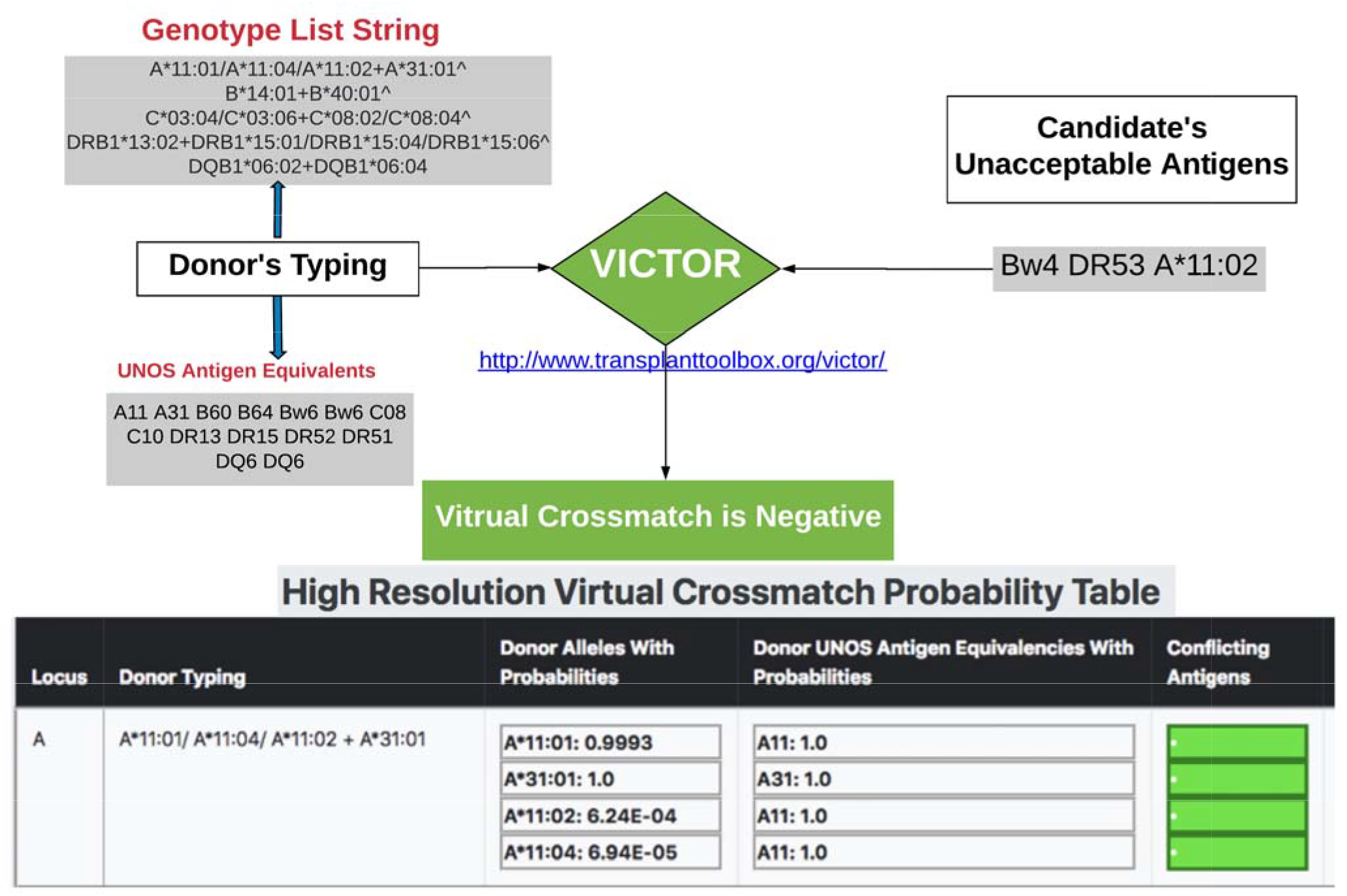
Illustration of a case where virtual crossmatch was computed negative by VICTOR when an unlisted allele-specific unacceptable HLA antigen had low risk probability

We also evaluated a similar counterfactual scenario to test what would have happened if the same candidate were listed with an allele-specific UA to A*11:01. Here we verified that if the VICTOR algorithm were integrated into the match run with a probability threshold, the donor would not have been offered as A*11:01 had >99% probability. We also found that this same donor would be offered by the current UNOS match run and would have to be turned down manually, needlessly increasing time-to-transplant. This case study illustrates that listing of two-field UAs would have far more utility if molecular typing data were captured and interpreted by UNet.

In a second case, a transplant candidate had a UA of DRB1*04:02 and was offered a donor typed with the following antigen: A2, A2, B51, B52, C03, C16, DR53, DR53, DR4, DR4, DQ7, DQ8. Manual analysis of the molecular typing report of the donor indicated that DRB1*04:02 was ruled out as a possible allele, so the offer was accepted. If provided a GL String, VICTOR could intermediately show that allele had been ruled out of the molecular typing, streamlining the VXM process. If the intermediate-resolution molecular typing data had been unavailable, a DR4 antigen-level typing would have provided a risk of 0.18 in VICTOR.

### Category #3 Cases: Refused offers where uncertainty in donor HLA typing revealed low DSA probabilities when interpreted by VICTOR

A candidate was listed with several UAs, including DRB1*12:02 and DRB1*13:03. An organ offer was made from a donor typed as DR12 and DR18. The molecular typing report was unavailable. The organ offer was refused, as there was lack of high resolution typing to determine if DRB1*12:02 was present in the donor. We analyzed this case with VICTOR by imputing the antigen-level typing and that risk probability was 0.80 if the donor was Asian / Pacific Islander, but less than 0.10 for every other race/ethnic category. In this case, either a molecular typing report or a VICTOR-based interpretation may have provided enough information to accept the offer.

A second case had candidate UAs DRB3*01:01 and DRB3*03:01 and a donor typing of DR52. While a molecular typing report was available, the offer was refused due to uncertainty as to if the UAs were DSAs. VICTOR provided risk probabilities of 0.36 for DRB3*01:01 and 0.11 for DRB3*03:01, which some centers may deem to be an acceptable risk and would proceed with a physical crossmatch.

## DISCUSSION

We built an HLA informatics tool to explore the benefits of having electronic utilization of molecular typing data in the solid organ allocation system. During the VXM, histocompatibility laboratories must manually interpret intermediate-resolution molecular HLA typing data when evaluating organ offers. Given an ambiguous donor HLA typing, many different alleles are possible. The probability of each allele can be assessed by interpreting the typing in the context of high-resolution population-specific HLA frequencies^32^. Because only two UNOS antigens per HLA locus can be represented in the UNet, typing ambiguity is not captured electronically. Current laboratory workflows for HLA typing of deceased donors usually cannot yield an unambiguous high-resolution typing in the time available. HLA informatics tools that can impute the probability of selected alleles or allele combinations can reduce uncertainty of the VXM in these situations. HaploStats is used by some centers, but it does not cross-reference UAs and does not return results for individuals with rare HLA alleles^25^.

To better inform the immunological risk from HLA typing ambiguity, we have built a tool to facilitate automated computation of VIrtual CrossmaTch with mOleculaR typing data (VICTOR). Our VICTOR tool compares the probabilities of all possible two-field IMGT/HLA alleles and UNOS antigens given the donor molecular typing and to a list of UAs for the transplant candidate.

We identified a category of offers made by the match run that were refused when the transplant center examined the more detailed donor molecular HLA typing report. Such attached reports are highly encouraged by UNOS but not yet mandated. In these cases, high risk of donor-recipient HLA incompatibility was fortunately identified by the transplant center before an incompatible organ would have been accepted and shipped. Here, a tool like VICTOR could speed HLA compatibility assessments and reduce the likelihood of errors. We envision that a standardized and automated interpretation of HLA typing data is feasible.

Because molecular HLA typing data is not systematically captured or utilized in UNet, we cannot easily speculate on the incidence rates of unexpected physical crossmatch that arise from HLA typing ambiguity or how often potentially-compatible offers are refused. Failure to interpret HLA typing data properly can result in adverse events such as unintended recipients and discarded organs. The utility of decision support systems such as VICTOR could be validated with prospective cohort-based study design that would include high-resolution typing and physical crossmatch. Positive physical crossmatch can occur for many reasons, making comparisons with VXM challenging to interpret as solid phase assays and physical crossmatch methods measure different antibody characteristics with differing levels of sensitivity and specificity^33^.

Our case studies of organ offers also illustrate how the allocation system might benefit from having a match run that could use typing data already in existence to facilitate more appropriate offers as opposed to vetting offers manually after allocation. If the match run could be programmed to avoid making offers with a high probability of donor-specific antibodies, the cold ischemia time and/or overall time-to-transplant could be shortened. Offers of incompatible donors could be reduced by setting either systemwide or center-specific thresholds for acceptable risk probabilities given the possible donor HLA specificities. Simulated allocation models that incorporate typing ambiguity and interpretation could assess the impact of these proposed changes on offer sequence length and acceptance rates.

The main purpose of our VXM tool is to provide an automated interpretation of ambiguous donor molecular typing data that would enable the collection of a standardized molecular typing data in Unet which would be used for virtual crossmatch decisions. There are many practical limitations to consider before this tool should be used for clinical decision making. Our initial implementation uses allele frequencies rather than haplotype frequencies so that a higher percentage of ambiguous typing can be interpreted than in HaploStats, however this increases the uncertainty in the probabilities provided^34^. Another category of limitations is related to the availability of reference HLA frequency information. High-resolution DQA1, DPA1, and DPB1 frequency data is not yet available because these loci were rarely typed in registry donors.

Calling of UAs from solid phase antibody assays carries an orthogonal set of challenges to interpreting typing data^35^. Antibodies are directed against epitopes that may be shared among many alleles within and across antigen categories^36^. There is substantial interest in performing a VXM based on mismatched amino acid motifs as putative epitopes^37^. Only the HLA antigens included in the current generation of solid phase assays are available in the current list of UNOS antigens in UNet. However, any IMGT/HLA allele can be listed as a UA in VICTOR. VICTOR relies on the transplant center to list any other less frequent alleles that may share putative antibody-reactive epitopes with the positive beads. Tools such as HLA Matchmaker can help determine which alleles to list as UAs^38^. The UNet system does yet not allow for amino acid motifs or “eplets” to be selected as UAs, though this is planned first for hypervariable region motifs of the DPB1 locus where serologic antigen categories were never adopted.

Discrepancy in HLA typing is one source of error for VXM, with an OPTN investigation finding that critical discrepancies that impact the match run are present in 2% of cases^39,40^. To reduce the incidence of HLA typing discrepancies, UNOS will soon require double-entry of keyed-in HLA data. However, this policy change will only address transcriptional errors. Electronic transmission of HLA typing data and automation of nomenclature translations could curb other categories of errors substantially. Towards this end, we published a complete mapping table between IPD-IMGT/HLA alleles and UNOS antigens as well as an ALLele-to-ANtigen (ALLAN) web tool^23^ to aid in deciding which antigens should be entered into UNet based on OPTN guidelines^24^. Manual keyed entry of ambiguous typing data would be very challenging due to the complexity of HLA data representations as fully-enumerated genotype lists or NMDP multiple allele codes. Electronic entry of molecular HLA typing data into UNet would be aided by adoption of data standards such as Histoimmunogenetics Markup Language (HML)^41^ by UNOS and typing kit vendors. Likewise, tools and data standards for antibody assay interpretation for entering UAs is an unmet need. The tool assumes that UAs were accurately determined from solid phase assays.

Retrospective outcomes studies that test hypotheses about the role of HLA matching in graft failure and development of de novo antibodies are hindered by the lack of molecular typing data for previous donors and recipients in the Scientific Registry for Transplant Recipients (SRTR) standard analysis files (SAF). HLA matching paradigms that assess longer-term immunologic risk are usually validated on more limited datasets where high resolution typing was available^42^. Imputation of antigen-level HLA data to amino acids has been performed on the SAF in a recent study of single amino acid mismatching and graft failure^37^. Systematic collection of molecular typing data would have an ancillary benefit of substantially reducing the uncertainty in the two-field HLA allele as well as amino acid assignments from imputation, which would improve the power for discovery^43^.

In this report, we provide a proof of concept tool to illustrate several advantages of a future match run implementation based on ambiguous molecular HLA typing data on top of existing antigen categories. VICTOR represents an important developmental step towards precision HLA compatibility assessments for solid organ allocation.

## ABBREVIATIONS

AFA: African American
API: Asian / Pacific Islander
ALLAN: ALLele to ANtigen
CAU: Caucasian
CPRA: Calculated Panel Reactive Antibodies
CWD: Common and Well-Documented Allele List
DSA: donor-specific antibodies
GL String: Genotype List String
HIS: Hispanic
HLA: human leukocyte antigen
HML: Histoimmunogenetics Markup Language
IMGT: International Immunogenetics Project
IPD: Immune Polymorphism Database
KAS: Kidney Allocation System
MAC: Multiple Allele Code
MFI: mean fluorescence intensity
NAM: Native American
NMDP: National Marrow Donor Program
OPTN: Organ Procurement and Transplantation Network
PyPI: Python Package Index
SAF: Standard Analysis File
SRTR: Scientific Registry of Transplant Recipients
UAs: unacceptable antigens
UNOS: United Network for Organ Sharing
VICTOR: VIrtual CrossmaTch for mOleculaR typing data
VXM: virtual crossmatch

